# Manipulating PARK7/DJ-1 Levels by Genotoxic Stress Alters Noncoding RNAs and Cellular Homeostasis

**DOI:** 10.1101/2025.02.03.636218

**Authors:** Keren Zohar, Haya Obied Zoubi, Michal Goldberg, Tsiona Eliyahu, Michal Linial

**Affiliations:** Department of Biological Chemistry, Institute of Life Sciences, The Hebrew University of Jerusalem, Jerusalem, Israel; Department of Genetics, Institute of Life Sciences, The Hebrew University of Jerusalem, Jerusalem, Israel

**Keywords:** Regulated cell death, Apoptosis, lncRNA, Pseudogene, Oxidation stress, X-ray, ribosome stability, miRNAs, RNA-seq, Parkinson diseases

## Abstract

DJ-1/PARK7 is a multifunctional protein that plays a vital role in sensing oxidative stress and maintaining redox homeostasis. As an oncogene, DJ-1 influences p53-mediated stress responses and contributes to cancer progression. This study investigates the impact of X-ray-induced DNA breaks on cellular responses post-irradiation under varying DJ-1 expression levels. Using siRNA knockdown and overexpression to modulate DJ-1 levels, transcriptional changes were analyzed through RNA-seq. Naïve cells exhibited only a moderate transcriptional response to X-ray exposure, including suppression of the cell cycle and activation of stress pathways. Overexpression of DJ-1 led to pronounced gene expression suppression, particularly suppression of most ribosomal and mitochondrial genes. Conversely, DJ-1 knockdown caused extensive, non-specific transcriptional changes, indicating disrupted cellular homeostasis. Notably, around 25% of non-coding RNAs were differentially expressed under these conditions. DJ-1 overexpression suppressed lncRNA of the host genes for snoRNAs, thus altering the miRNA sponging capacity after X-ray exposure. These findings underscore DJ-1’s critical role in modulating cellular responses to genotoxic stress, reshaping transcriptional landscapes, and regulating non-coding RNA profiles. DJ-1 emerges as essential for maintaining genomic stability.

## Introduction

PARK7 (DJ-1) is a multifunctional protein that has been implicated in a variety of human diseases (1–3). DJ-1 is expressed in cells and tissues to scavenge reactive oxygen species (ROS). Furthermore, it acts as a chaperone, regulates transcription, and is involved in signal transduction and mitophagy (4). DJ-1 was initially identified as an oncogene and was later discovered as a causative gene for Parkinson’s disease (PD) (5), and other age-related diseases (6). Different aspects of PD pathologies, including cell death, α-synuclein aggregation, and the formation of amyloid-like filaments, were attributed to mutations in the PARK7 gene (6). In addition to the direct role of DJ-1 in protecting neurons from oxidative damage, DJ-1 was implicated in inflammatory (3) and metabolic diseases (5). Similar to the impact of mutations in DJ-1 on cell death and PD development, depletion of DJ-1 in pancreatic beta cells reduced the capacity of the cells to cope with stress conditions, and make them prone to diabetes (7). Manipulations of DJ-1 in animal models confirmed its causal role in obesity-induced inflammation (8), metabolic reprogramming, energy homeostasis (9) and other physiological conditions (10).

At the molecular and cellular levels, the human PARK7 gene encodes DJ-1, a ∼20 kDa protein that forms a dimer for its function. The homodimer is predominantly localized to the mitochondria, where it plays a role in the regulation of electron transport, apoptosis, and mitophagy (11). As a sensor for oxidative stress, it protects cells from damage by ROS (4,12) and its cellular activity is regulated by the oxidation status of key cysteine residues (13). Upon oxidative stress, DJ-1 scavenges ROS and acts in the mitochondrial quality control machinery (14). Its role in protecting neurons is related to DJ-1’s function as a major antioxidant (15). Specifically, via the Keap1-Nrf2 pathway that governs the redox homeostasis within cells (16). Under oxidative stress, DJ-1 activates and stabilizes Nrf2 by dissociating it from Keap1 and preventing its ubiquitination. By promoting the entry of the Nrf2 transcription factor (TF) to the nucleus, it can activate the expression of numerous antioxidant and detoxifying genes. When DJ-1 is depleted, antioxidant enzymes are diminished, and apoptosis and cell death are induced (17).

In addition to the substantiated role of DJ-1 in the oxidative stress response, multiple studies show a role for DJ-1 as a mitogen-dependent oncogene that promotes the progression of various cancers. High expression of DJ-1 was linked to poor prognosis in numerous cancer types (e.g., brain tumors, breast cancer, non-small cell lung carcinoma, prostate cancer, pancreatic carcinoma, hepatocellular carcinoma, colorectal cancer, and more). In most cases, elevated expression correlates with aggressive disease, reduced survival, and metastasis (18). Several mechanisms have been proposed in which DJ-1 supports tumor progression. For example, stable interaction of DJ-1 with anti-apoptotic factors (e.g., Bcl-Xl) will prevent the release of mitochondrial factors such as cytochrome c and apoptosis-inducing factor (AIF) into the cytoplasm (19). When such apoptosis-inducing factors accumulate in the cytoplasm, they directly activate the caspase cascade and induce apoptosis. A functional link between the roles of p53 and DJ-1 was proposed in the context of oxidative stress but also via its nuclear function. Specifically, a feedback loop in which DJ-1 mediates the activation of p53 in response to oxidative stress. It was also suggested that the loss of p53 leads to increased DJ-1 expression in transformed cells, which is signified by oncogenic activation of AKT. DJ-1 overexpression is also associated with its ability to inhibit tumor suppressor proteins like PTEN. By inhibiting PTEN, PI3K/AKT signaling pathway is activated, promoting cell growth and survival, which govern the poor prognosis in these cancers (20).

The response of DJ-1 to oxidative stress and to elevated ROS has been established. In addition to a physiological formation of ROS, X-ray irradiation also generates ROS, such as hydroxyl radicals and hydrogen peroxide, which, at high levels, cause damage to DNA, lipids, and proteins. Whether DJ-1 is part of the DNA damage response (DDR) remains unknown. Following X-ray irradiation, cells effectively activate the DDR, which is a network of processes that includes sensing double-strand breaks (DSBs), and activate repair mechanisms (21). The activation of DDR primarily arrests replication of the damaged DNA to allow the activity of DNA repair mechanisms such as homologous recombination (HR) and non-homologous end joining (NHEJ). In cases where the damage is too severe, cells will eventually die by programmed cell death, senescence, or other forms of regulated cell death (22). A recent study suggested that in the context of PD patients, the pathological mutation of DJ-1 impairs the activity of DNA repair function. This is in agreement with observations from DJ-1 knockout mice that show an increase in DSBs and genome instability. The notion is that unrepaired DNA damage may contribute to PD pathology (23). Along the PD progression, neurons with defective DNA repair will drive genome instability, mitochondrial dysfunction, and eventually neurodegeneration.

In this study, we asked whether we can detect a cell response and a shift in cell decisions following X-ray irradiation under controlled conditions where the DJ-1 expression levels are molecularly manipulated. Specifically, we tested the impact of changing the DJ-1 levels from their baseline levels to cells with knockdown (KD) and overexpression (OX) of DJ-1 and monitored their overall response. We identified global changes in transcription as well as consistent changes in the amount and diversity of ncRNAs and lncRNAs in particular. Changes in cellular homeostasis, regulated cell death, and translation were revealed in cells challenged with X-ray using varying levels of DJ-1 as a baseline.

## Materials and methods

### Cell viability using flow cytometry (FACS)

Human embryonic kidney cells 293 (HEK293 from ATCC) were cultured in 6-well plates using Dulbecco’s modified Eagle’s medium high-glucose with 10% fetal bovine serum to reach a 70- 80% confluence level at 37°C and 5% CO2 before analysis by fluorescence-activated single-cell sorting (FACS). HEK293 cells were incubated with the fluorescent dye propidium iodide (PI) and Annexin V- FITC, purchased from MBL (MEBCYTO-Apoptosis Kit; Woburn, MA, USA). Cells were stained with Annexin V according to the manufacturer’s protocol from MBL. The fluorescence of Annexin V indicates the presence of phosphatidylserine at the outer leaflet of the plasma membrane. Apoptosis and other phases of cell death were discriminated by staining with PI and Annexin V (24,25). After staining, live cells show little or no fluorescence (Annexin V-/PI-), while early apoptotic cells show FITC fluorescence (Annexin V+/PI-). The late apoptotic (and necrotic) cells display red and green fluorescence (Annexin V+/PI+) (26). FACS analysis was conducted with 30,000 cells, and the fraction of dead cells stained with PI was determined.

### Knockdown by siRNAs

Cells were cultured to reach a 20–30% confluent level (3×10^4^ per cm^2^). For transfection, we used Lipofectamine 2000 (Lipo2000; Invitrogen, Cat # 11668019, Carlsbad, CA, USA) in cells at 40–50% confluency according to the manufacturer’s instructions. Lipo2000 was shown to be ideal for HEK293 (27,28). The esiRNA (Mission system, Sigma-Aldrich, Burlington, MA, USA) consists of a pool of hundreds of siRNA (21 bp each), where each individual dsRNA has a low concentration in the pool, which diminishes most off-target effects, while producing an efficient knockdown. We apply the esiRNA-RULC of a mixture of 21 nt dsRNA used as a negative control (RLUC stands esiRNA directed against Renilla Luciferase; EHURLUC) in addition to the specific PARK7 (to knockdown the DJ-1 transcripts and protein products; EHU113961). RULC experiments were used to measure the baseline effects of introducing esiRNA that controls for the delivery method and allows to distinguish the cellular response to the targeted siRNA itself (29). After waiting for 18 hours cells were exposed to X-ray radiation (Parameters: 12.5mA, 320KV, 1000cGY). After waiting for an additional 6 hours, RNA extraction was done.

### Overexpression of DJ-1

One day prior to the transfection, the cells were plated in a 10-cm plate at an initial density of 10× 10^6 cells per plate. The expression of pCMV3 vector for the canonical DJ-1, fused to doble Strep-tags at his C-terminal by Sino Biological company. The Strep-tag peptide exhibits an intrinsic high affinity towards Strep-Tactin engineered streptavidin. Expression plasmid (DJ-1/empty vector of pCMV3) and PEI (260008-5; Polysciences) were separately diluted in Opti-MEM1 (31985- 047; Gibco) and mixtures were incubated at room temperature for 25 min to allow polyplex formation prior to its addition to the cell culture. The mixture was added dropwise onto the cells. After waiting for 18 hours cells were exposed to X-ray radiation (Parameters: 12.5mA, 320KV, 1000cGY). After waiting for an additional 6 hours, RNA extraction was done.

### Western Blot

Cells were lysed using 2xSDS with DTT lysis buffer, incubated at 95°C for 5 min, and frozen at −20°C. Before use, lysates were re-incubated at 95°C for 5 min followed by a short spindown to remove debris. Protein lysates were electrophoresed on 13% SDS polyacrylamide gels followed by semi dry transfer to nitrocellulose membranes that were blocked with TBS-T buffer (0.15 M NaCl, 0.05 M Tris hydroxymethyl methylamine and 0.1% Tween-20, pH 7.4–7.6) containing 5% skim dry milk (BD Diffco) for 0.5 h at room temperature. Membranes were then washed with a TBS-T buffer and incubated overnight at 4°C with primary antibodies, diluted according to manufacturer instruction in 4% BSA. Membranes were washed with TBS-T buffer and incubated with secondary antibody (1:1000 in TBS-T with 4% skin milk) for 1 h at room temperature, followed by three washes with TBS-T, 10 min each. Chemiluminescence was visualized with an ECL detection kit (Biogate, Ness Ziona, Israel) using gel imager (ChemiDoc, Bio Rad, Hercules, CA, USA or Fusion FX, Viber Lumart, Marne-la-Vallée, France) and quantified using the software program Image Lab 6.1. Results were normalized to loading control (Tubulin).

### Reverse Transcription PCR (RT-PCR) and PCR

Cells were collected for RNA preparation. Total RNA was extracted from cell culture with TRIzol (Thermo-Fisher Scientific, Waltham, MA, USA), and RT was performed using a Ready-To-Go first-strand synthesis kit (Cytiva, Marlborough, MA, USA) according to the manufacturer’s instructions. RNA was reverse transcribed into cDNA (1 μg) and used in the PCR reaction. The PCR conditions consisted of denaturation at 95°C for 2 min and 35 cycles (10 s at 95 °C, 15 s at 60°C, 30 s at 72°C), and 5 min for a final extension. The PCR products were separated on 1.5% agarose gel and stained with ethidium bromide, followed by densitometry measurement (using image processing ImageJ program Ver 1.54, from GitHub). The primers were designed against human RefSeq by the Primer3Web tool (Ver 4.1.0). The forward (F) and reverse (R) primers of β-actin (196 nt) were F: CATGCCCACCATCAGCCCTGG and R: ACAGAGCCTCGCCTTTGCCGA, respectively. For DJ-1 (376 nt), the primers were F: GCCTGGTGTGGGGCTTGTAA and R: GCCAACAGAGCAGTAGGACC. For DJ-1, which is only suitable for transcripts from the plasmid (and not endogenous) (447 nt), the primers were F: CAGTGTAGCCGTGATGTGGT and R: AGCAGACCCCGCGTCTTTA.

### RNA-Seq

Total RNA was extracted using the RNeasy Plus Universal Mini Kit (QIAGEN, Cat # 73404, Redwood City, CA, USA), according to the manufacturer’s protocol. Total RNA samples (1 μg RNA) were enriched for mRNAs by pull-down of poly(A). Libraries were prepared using a KAPA Stranded mRNA-Seq Kit, according to the manufacturer’s protocol, and sequenced using Illumina NextSeq 500 to generate 85 bp single-end reads (a total of 25–30 million mapped reads per sample).

### Bioinformatic Analysis

Next-generation sequencing data underwent quality control using FastQC, version 0.11.9. They were then preprocessed using Trimmomatic (Ver. 0.32) and aligned to the reference genome GRCm38 with the STAR aligner (Ver. 2.7.0a) using default parameters. Data was normalized to account for data inflation and converted to TPM (transcripts per millions) to corrects for gene length. TPM values are used for comparing between genes. Genomic loci were annotated using GENCODE (release 46). Genes with low expression were filtered out of the dataset by setting a threshold of a minimum of two counts per million in three samples.

Differential analyses were performed on all experimental groups, and genes with an FDR < 0.05 were considered. Only genes with an absolute log fold change of ≥0.5 were labeled up- or downregulated, all the rest were considered unchanged. We partition genes by type as coding and non-coding (including pseudogene, anti-sense, miRNAs, TEC, lncRNA, and other rare biotypes). Due to the overexpression of PARK7 (expressed for 4.5% of all normalized reads, we used a conservative view where we removed the expression of PARK7 and recalibrated the data prior to differential gene expression (DEG) analysis.

RNA-seq experiment results are visualized by an MA plot that transforms the data into M (log ratio) and A (mean average) scales. The functional analysis and network view was based on STRING protein– protein interaction (PPI) platform. The minimal PPI confidence score ranges from 0.7 to 0.9. We used the gene expression normalization nTPM as the basal expression of DJ-1 in HEK293 as reported in Human Proteome Atlas (HPA).

For functional enrichment of ncRNAs we applied the analysis of LncSEA (Ver 2.0). This tool uses that uses predetermined sets of lncRNA functions supported by 400k reference lncRNA sets involved in main 33 annotated categories (e.g., chromatin, TF, RNA-RNA interaction, cell cycle and more) that are used for class enrichment analysis (30). Data of lncRNA sets are based on integrating of TF ChIP-seq, DNase-seq, ATAC-seq and H3K27ac ChIP-seq data associated with hundreds of human cell types (30). The analysis of the Simpson index assess reflects the diversity in the functional roles of lncRNAs in LncSEA2.0 database. A lower index indicates a concentration of lncRNAs in fewer functions. For RNA-protein interaction we used the compilation and unification of results from RNAInter4.0, NPInter 4.0 and ENCORI (supported by lncSEA2.0). For an indirect connectivity map lncRNAs, genes and miRNAs are connected by the lncTAR 2.0 and ncPATH tools. The tool integrates approximately 32k verified lncRNA–target interactions, 5k experimentally validated ceRNA networks and 0.5M potential connection to coding genes, across 220 KEGG pathways (31).

### Statistical Analysis

Principal component analysis (PCA) was performed using the R-base function prcomp (R studio Ver. 4.1.0). EdgeR (Ver. 3.36.0) was used to perform RNA read counts by the trimmed mean of the M-values normalization of RNA (Trimmed Mean of M-values; TMM) and differential expression analysis. Figures were generated using the ggplot2 R package (Ver.3.3.5).

## Results

### DNA damage response by X-ray irradiation

To test the impact of X-ray on cell decisions with varying levels of DJ-1 in cells, we optimized the experimental conditions to achieve an efficient DDR following X-ray while minimizing cell death. The phosphorylation of histone H2AX on Ser139 (γ-H2AX) is a transient marker for a local DDR at the site of DSBs (32). A mock experiment, involving the overexpression of an empty plasmid, confirmed that the maximal signal of γ-H2AX is observed 1 hour after irradiation, where the level of modified H2AX declines post-X-ray irradiation (Supplemental **Figure S1**). **Figure 1A** shows the impact of DJ-1 manipulation through overexpression (OX). We compared the endogenous protein levels of cells before and after X-ray irradiation using mock transfection (empty plasmid), and cells that were overexpressed with DJ-1. Two bands are detected for the OX conditions: the endogenous DJ-1 protein (24 kDa) and a 27 kDa band that represents the double Strep-tag protein product (see Methods). The strong signal of the OX DJ-1 has no effect on the endogenous protein level. The levels of endogenous DJ-1 remain stable across all conditions, and are unaltered by irradiation (6 hours post 10 Gy X-ray). We then tested whether the molecularly manipulated cells cope with X-ray irradiation by monitoring cell viability. Testing naïve untreated cells (N.T.) revealed that the fraction of dead cells is minimal (4%), with a moderate decline in cell viability only 24 hours after irradiation (Supplementary **Figure S1**), and no increase in apoptosis during this time window (Supplementary **Figure S1**). We concluded that while DDR was coordinately activated immediately after the DNA damage (as shown by γH2AX) at 6 hours post-irradiation, the long-term effects of apoptosis and cell death were minimal.

**Figure 1.**
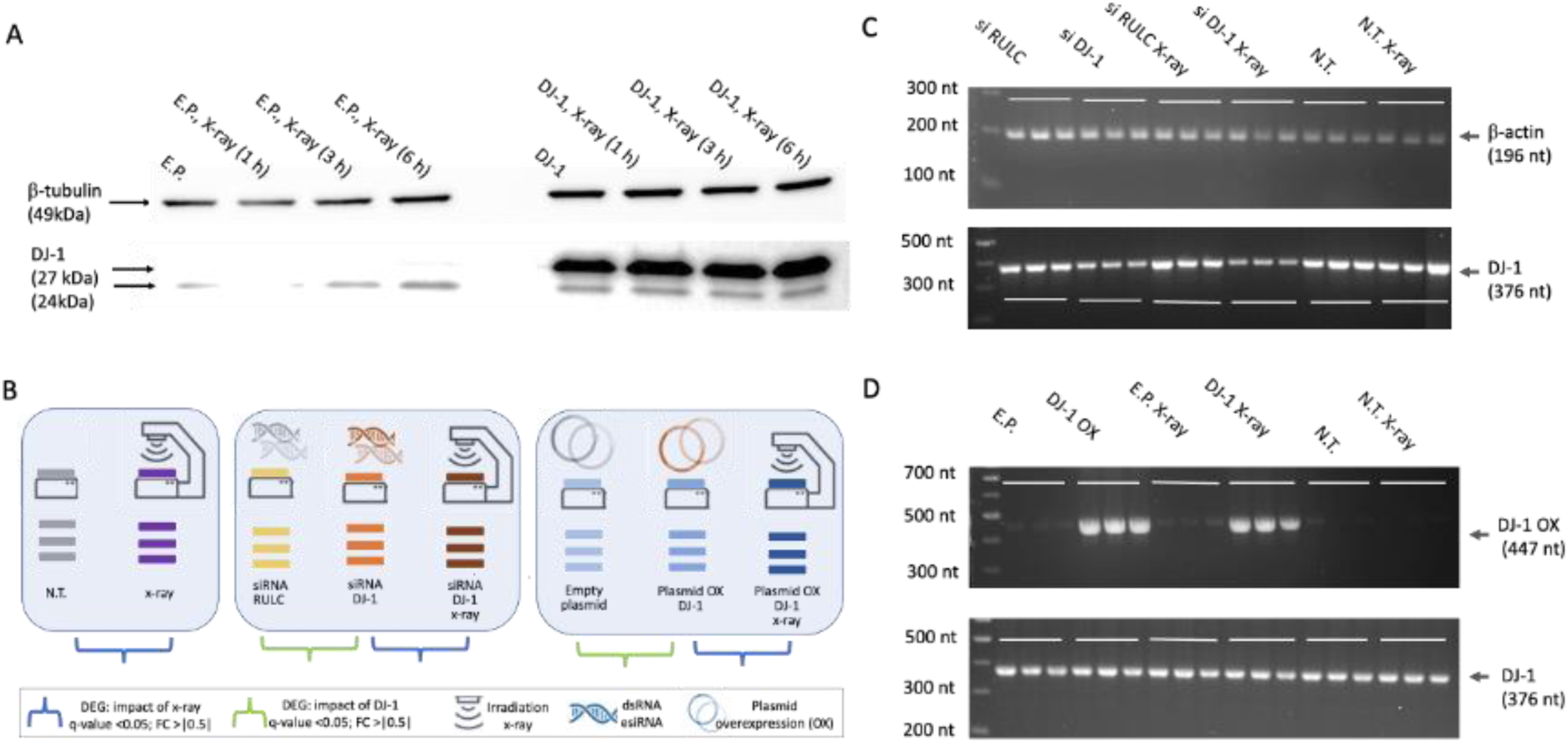
The outline of the comparative RNA-seq experiment in HEK293 cells with different baseline levels of DJ-1 following X-ray (10 Gy, 6h post irradiation). (A) Western analyses of the cells following X-ray treatments for 1-6 hours post irradiation for a mock experiment (empty plasmid, left) and DJ-1 tagged plasmid (right). (-tubulin is used as an internal control. (B) Schematic illustrations of the sets of experiments used for the RNA-seq data. Each experimental group is composed of biological and technical triplicates with and without X-ray exposure (10 Gy). Pairs of experimental groups are marked by blue and green parentheses. The blue pairs will be discussed (total of 6 conditions, 18 RNA-seq sequenced libraries). DEGs are defined as genes that passed the pre-determined thresholds of log2(FC) >|0.5| and adjusted p-value FDR <0.05. (C) RT-PCR products of treated cells for the indicated conditions are shown. The PCR products were separated in agarose gel (2.5%) for the amplicons of DJ-1 and (-actin as an internal control. Results from the untreated (N.T.) and DJ-1 knockdown (KD with siRNA) are shown. (D) RT-PCR results for the indicated conditions for DJ-1 overexpression (OX, transfected with DJ-1 plasmid) and N.T. Top: RT-PCR using oligonucleotide primers to detect overexpressed DJ-(DJ-1) plasmid. Bottom: Detecting endogenous DJ-1 levels in cells using primers specific to the endogenous DJ-1 (see Methods).

### Testing the impact of X-ray irradiation in cells ranges by different levels of DJ-1

Exposing living cells to X- ray indices double-strand breaks (DSBs) and, as a byproduct, an elevation of oxidative stress. We anticipate that the presence of DJ-1 is linked to the cellular response at varying redox levels. To study the transcriptomic signature of cultured cells following X-ray irradiation, we manipulated the baseline levels of DJ-1 through knockdown (KD) and overexpression (OX) protocols. **Figure 1B** depicts the overall design of the experiment in which cells were irradiated with X- rays while the expression level of DJ-1 was altered through molecular manipulation. In this study, we focus on the impact of X- ray treatment on cultured cells (HEK293) under naïve conditions (N.T.), in cells where DJ-1 expression was suppressed using the siRNA approach (denoted DJ-1 KD), and in cells with DJ-1 overexpression (DJ-1 OX). Each experiment was conducted under identical conditions to compare the effects of short-term irradiation (6 hours post-10 Gy). All experiments were normalized (using TMM; see Methods). Here, we report on the transcripts (coding and non-coding genes) that were significantly altered in response to X-ray genotoxic stress.

Figure 1C shows the results from RT-PCR of cells collected 6 hours after exposure to X-ray while monitoring the endogenous and manipulated levels of DJ-1 in the different cellular settings along with the appropriate controls. Figure 1C shows that only the KD of DJ-1 caused a suppression in the DJ-1 transcript, but siRNA for non-specific sequences (siRNA RULC, see Methods) had no effect on the basal DJ-1 level. Similarly, there was no detectable difference in naïve conditions (N.T.) or following X-ray for DJ-1 transcripts. We asked whether the siRNA protocol with DJ-1 resulted also in suppression of the DJ-1 protein. Supplementary **Figure S2** shows the results of a Western blot following the knockdown protocol. The effect on the DJ-1 protein was evident for DJ-1 KD but not for the non-specific siRNA RULC (see Methods). We concluded that the KD protocol is specific and reduced the original protein level to negligible level (Supplementary **Figure S2).** To ensure the specificity of the RT-PCR, we used oligonucleotide primers that can only detect sequences specific to the introduced DJ-1 plasmid (Figure 1D). The results show a strong signal for DJ-1 OX relative to cells transfected with an empty plasmid. Notably, despite the addition of an OX plasmid, the level of endogenous DJ-1 remained stable, as confirmed for the protein levels (Figure 1A).

### Global statistics of transcriptomes of cells exposed to X-ray

Figure 2A shows the changes in the transcriptome by exposing cells to X-ray irradiation with statistics for the significant differentially expressed genes (DEG) from RNA-seq experiment. The normalized level of endogenous DJ-1 and following KD protocol are shown. The DJ-1 KD reduced the original transcript level by >8 folds (33) with no effect on reduction in the basal DJ-1 levels using siRNA RULC (non- specific; see Methods). Figure 2B shows the results of all 9 samples for the DJ-1 OX setting. The OX DJ- 1 but not the control of an empty plasmid resulted is a drastic elevation in DJ-1 transcripts by at least 200 folds. To test the quality of the RNA-seq data, we analyzed the normalized sequenced reads using principal component analysis (PCA, Supplemental **Figure S3)**. Approximately 44% and 49% of the variance was explained using the PC1 and PC2 for DJ-1 OX and the DJ-1 KD experiments, respectively (Supplemental **Figure S3**). We concluded that the experiment is robust with reliable clustering of the biological triplicates for the different experimental groups.

**Figure 2.**
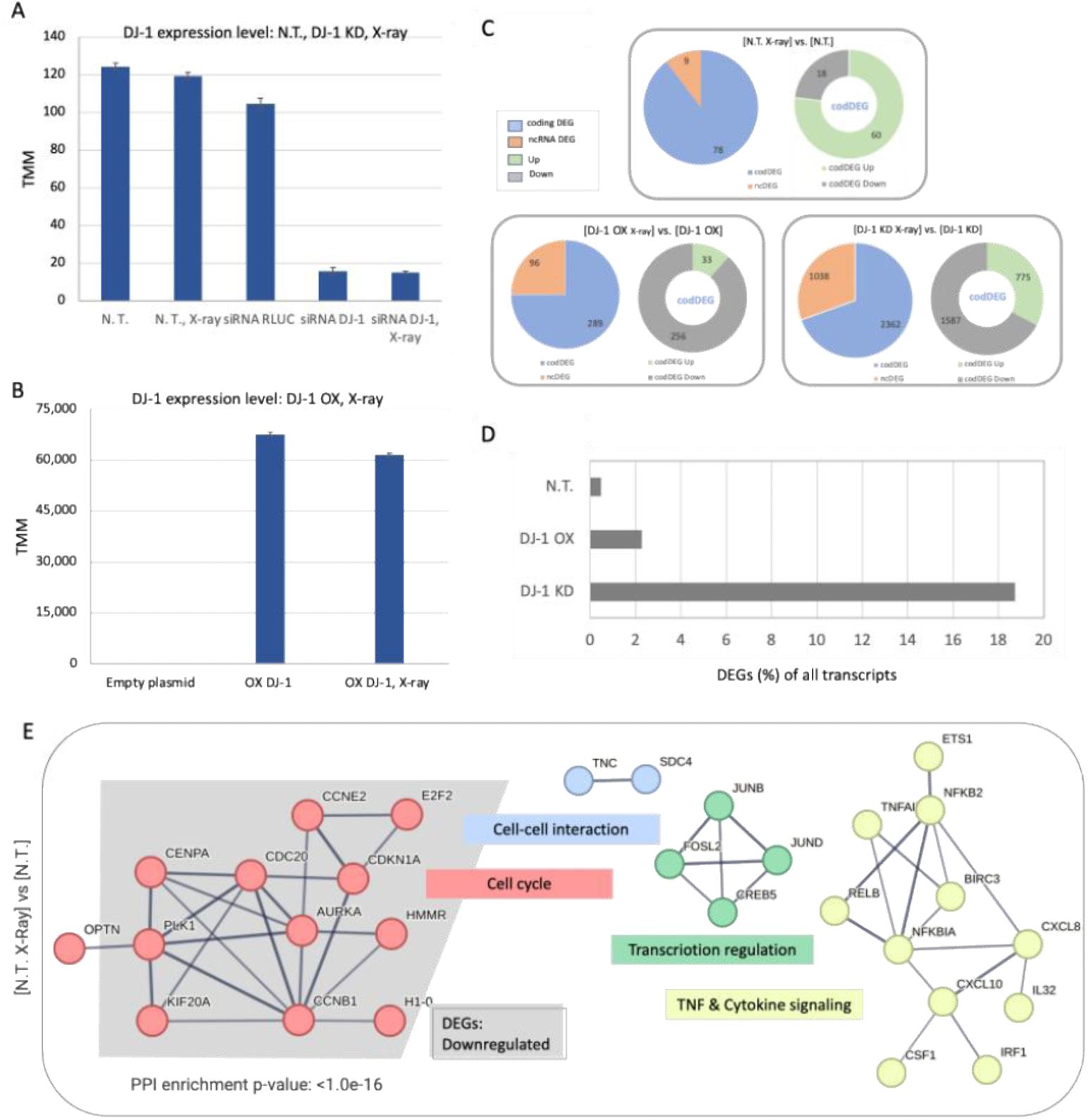
Transcriptome view of cell response to X-ray irradiation. **(A)** Expression of DJ-1 transcript across the cellular settings of N.T., DJ-1 KD with and without X-ray irradiation and siRNA RULC as non-specific control. The level of expression is according to TMM normalization. **(B)** Expression of DJ-1 transcript across the cellular settings of DJ-1 OX with and without X-ray irradiation. A control of empty plasmid is included. **(C)** A summary of the DEG statistics for prior and post X-ray irradiation for all three cell settings: N.T., DJ-1 OX and DJ-1 KD. Significant DEGs are marked as Up- and Downregulated (log_2_(FC) >|0.5|) for all 3 experimental settings. For each setting, the DEGs are compared relative to their matched control. Each analysis shows the number of DEGs according to coding (codDEG, blue) and noncoding RNAs (ncDEG, orange). Partition of the DEGs to those that are upregulated (Up, green) and downregulated (Down, gray) is shown. N.T., non-treated cells. **(D)** The fraction (%) of DEGs from all identified transcripts for N.T. and cells where the DJ-1 levels were manipulated (marked DJ-1 OX and DJ-1 KD). **(E)** STRING PPI network for codDEG of N.T. X-ray vs N.T. Only the subnetworks with ≥2 interacting genes are shown (p-value: <1.1 e-16; STRING confidence score >0.7). The function associated with each subnetwork is color coded. The gray background in the Cell cycle network marks the downregulated genes.

Comparing the extent of DEG in N.T., DJ-1 OX and DJ-1 KD experiments showed that the stronger overall response was for DJ-1 KD, with 3400 affected genes, among them 30% are noncoding DEG (ncDEG) (Figure 2C). In contract, for the N.T. setting, the fraction of ncDEG is only 10% after X-ray irradiation, and the fraction of ncDEG in response to X-ray is 25% for the OX setting. The increase in the fraction of ncDEG in the DJ-1 OX and DJ-1 KD settings are statistically significant (hypergeometric distribution p-value: 0.003 and 7.7e−29, respectively). Focusing on ncDEG, we observed 10 times more genes in DJ-1 KD relative to DJ-1 OX setting (1038 relative to 96) and 100 folds more relative to N.T. cells (9 ncDEG) (Figure 2C). We concluded that representation of ncDEG varies greatly with respect to the state of DJ-1 in cells. We also observed that following exposure to X-ray the majority of the coding DEGs (codDEG) are associated with downregulation (Down, gray). For the DJ-1 OX the upregulated genes (Up, green) accounts of only 11% of the codDEG (33 genes). Supplementary **Table S1** lists all codDEG and ncDEG along with the statistically significant and the normalized fold change in expression.

For testing the impact of the X-ray irradiation on the transcriptional signature, we indicated the fraction of DEGs (coding and non-coding genes) relative to the background control level. Figure 2D specifies the fraction of DEGs in responded to X-ray under different DJ-1 cellular levels. While only 0.5% of the identified genes are considered DEGs for the N.T., cells depleted from DJ-1 (KD) resulted in an extended and stronger response (>18% DEGs).

### Naïve cells display a limited but coordinated response to X-ray irradiation

Figure 2E shows protein-protein interaction (PPI) network of the 78 codDEG from the N.T. setting (STRING-based p-value: <1.1 e-16; confidence score ≥0.7). Most genes (60) were upregulated and only 18 were downregulated (Figure 2C). Notably, the network of cell cycle enriched network is almost entirely composed of downregulated DEGs (Figure 2E, gray background). The other subnetworks of TNF signaling, transcription and cell-cell interaction include the upregulated (Up) DEGs. The TNF signaling network is centered around RelB, NFKBIA and NFKB2 that are interact with each other to form distinct transcriptionally active complexes that together with NF-κB orchestrate cell survival pathway (Figure 2E). NF-κB is a pleiotropic transcription factor (TF) involved in processes such as inflammation, differentiation, cell growth, tumorigenesis and apoptosis. Thus, to overcome the genotoxic effect caused by X-ray on cell viability, numerous protection processes were activated along a coordinated suppression of cell cycle.

### Downregulation in ribosomal and mitochondrial transcripts dominates DJ-1 overexpression

The transcriptomes from cells with a baseline level of DJ-1 (N.T.), overexpressed and depleted levels of DJ-1 were compared according to the trend of upregulated (Up, Figure 3A) and downregulated (Down, Figure 3B). We observed that the number of overlapping codDEG across the different cellular settings is limited. While the union of all codDEG in >2800, the number of genes at any intersection is negligible (2%), arguing for unique transcriptional program is induced for each setting (Supplementary **Table S1**). An unexpected observation concerns the exceptionally high number of codDEG in DJ-1 KD (2362 codDEG). Despite the very high number of DEGs, a negligible number of them overlap with the other two experimental settings. This observation is statistically significant (hypergeometric distribution test p-value: 6e-05). Still, the 11 genes are shared between N.T., DJ-1 OX and DJ-1 KD (Figure 3C, green frame) belong to the network of TNF and NF-kB signaling. Additionally, the 13 DEGs overlap between the N.T. and DJ-1 OX cells (Figure 3D, yellow frame) forms a functional cell division enriched network. We concluded that in all cellular settings following X-ray irradiation, cell division is suppressed and TNF and NF-kB signaling pathways are activated.

**Figure 3.**
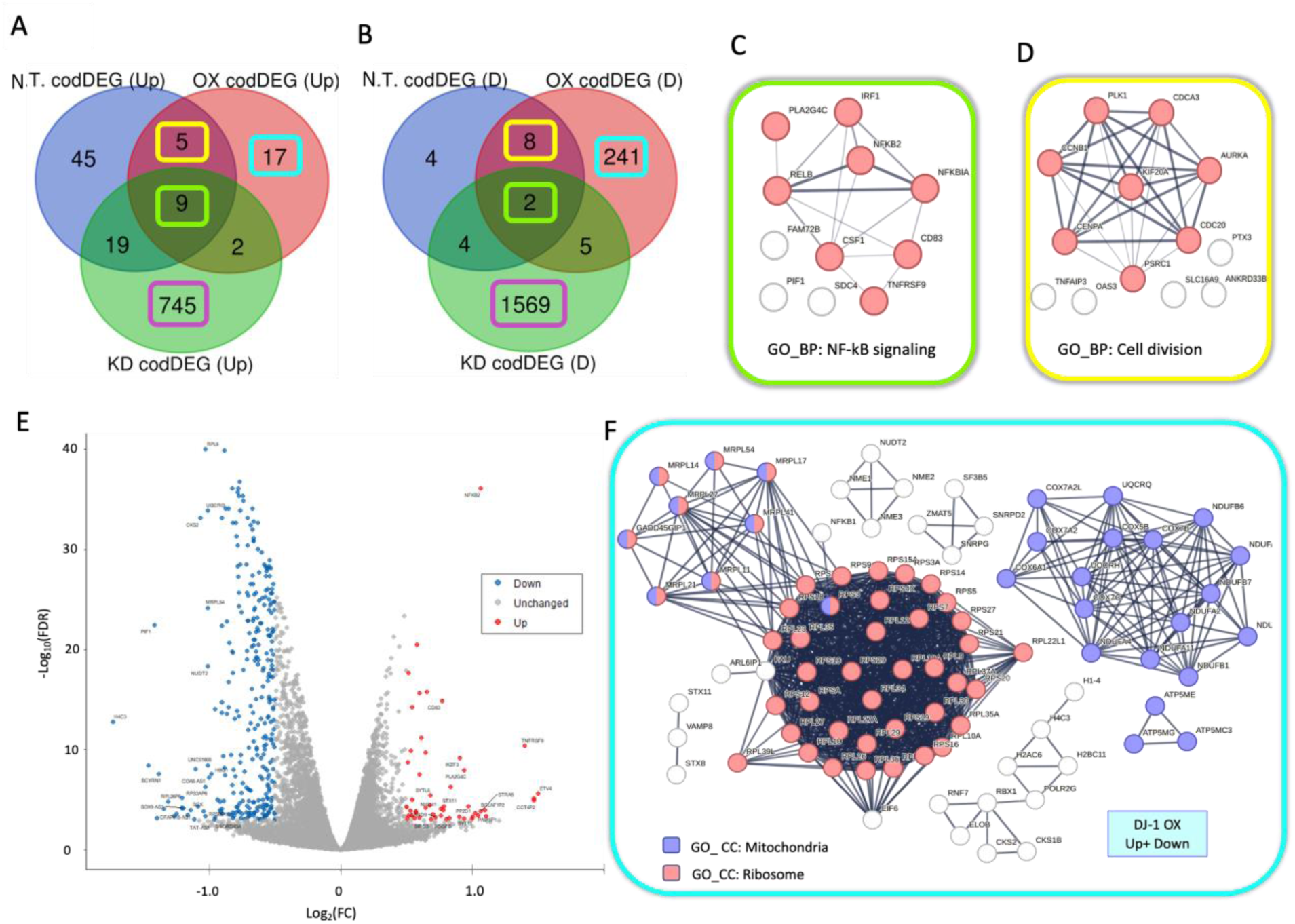
Analyzing the codDEG for N.T., DJ-1 OX, and DJ-1 KD. **(A)** Venn diagram of the upregulated codDEG (marked Up). **(B)** Venn diagram of the downregulated codDEG (marked Down). The shared genes between all three cellular settings are indicated by light green frame. Additional codDEG that overlap the N.T. and DJ-1 OX are colored by a yellow frame. The unique DJ-1 OX set is colored by light blue frame. **(C)** STRING-based PPI networks (confidence score ≥0.7) for the 11 shared DEGs by the green colored frames shown in **Figures 3A-3B**. **(D)** STRING-based PPI networks (confidence score ≥0.7) for 13 DEGs, marked by the yellow-colored frames, shown in **Figures 3A-3B**. **(E)** Volcano plot of the DJ-1 OX X-ray vs cells with DJ-1 OX. The plot shows the fold change (FC) by log_2_(FC) and minus log10(FDR). **(F)** STRING-based PPI networks (confidence score ≥0.9) for the 258 unique DJ-1 OX DEGs by the blue colored frames shown in **Figures 3A-3B**. Only networks of ≥3 genes are shown. The colored nodes are annotated by the colored frame for the matched gene ontology cellular component (GO_CC) annotation. The connectivity of the STRING results for the shared genes is statistically significant (p-value <1.e- 16). The codDEG are listed in Supplementary **Table S1**.

Figure 3E shows the results by a Volcano plot of the DJ-1 OX X-ray vs. control cells of DJ-1 OX. Most genes were strongly downregulated (blue dots). However, a few significant upregulated DEG belong to NF-kB signaling (as in Figure 3C). We then tested the nature of the unique codDEG (17 upregulated and 241 downregulated genes; Figures 3A**-3B**) for the DJ-1 OX setting. Analyzing these codDEG (Figure 3F, light blue frame) identified a large fraction of almost all ribosomal protein genes (Figure 3F, red nodes). The network is extended to the genes of the mitochondrial ribosomes (Figure 3F, nodes colored purple/red). In addition, a tight network of downregulated DEG (Figure 3F, purple nodes) are mitochondrial energy production genes. Additional subsets comprised from downregulated genes include a network of chromatin modifying genes (10 genes, H2AC6, RBX1 as representatives) and smaller subnetworks with a few DEGs acting in secretion (e.g., VAMP8), splicing machinery (e.g., SF3B5) and NME family members (*e.g.,* NME1-3). The NME genes carry a wide range of cellular functions, such as energy metabolism, cytoskeleton dynamics and DNA repair. For example, the NME1 is recruited to sites of DNA damage, acts in the proofreading and specifically in determine the mode of DSB repair (34). We concluded that 6 hours after X-ray induction, DJ-1 overexpression in cells triggered a coordinated transcriptional wave, leadin to the suppression of major cellular processes, including chromatin-transcription axis, energy production and primarily a suppression of the translation machinery.

Analysis of the codDEG from DJ-1 KD X-ray relative to DJ-1 KD identified large number of significant genes (>2300 genes). Despite the large number of codDEG the PPI of the STRING based network was rather limited (Supplementary **Figure S4**). Moreover, only very broad functional enrichment was detected. Functional enrichment of the upregulated gene set (745 genes, Figure 3A) identified cell junction and membrane, while for the downregulated codDEG it was mainly nuclear functions and transcription regulation (Supplementary **Figure S4**). We postulate cells depleted from DJ-1 lack robustness and displayed increased cells’ fragility to X-ray irradiation. For the DJ-1 KD condition, the cells display a global change in nuclear function associated with a global suppression of transcription and alteration of chromatin. Functional enrichment analysis for codDEG for the setting of DJ-1 KD X-ray relative to DJ-1 KD is accessible in Supplementary **Table S2** (partition to up and downregulated codDEG).

### Induction of ncRNAs in response to X-ray irradiation

The RNA-seq results show that about 19% of all transcripts are classified as ncRNAs (totaling 16,825 genes filtered by a minimal expression threshold; see Methods). A significant observation indicates that the expression levels of ncRNAs are, in general, very low (Supplementary **Figure S5**). Specifically, only 3.4% of the total expression levels are associated with ncRNAs (3,230 genes), while the remaining (96.6%) are associated with coding genes. About 75% of the ncRNAs exhibit low expression (<2 TMM), 8% of the ncRNAs are expressed with ≥10 TMM, and only 14 ncRNAs are highly expressed (≥100 TMM, Supplementary **Figure S5**). Partitioning of the 3230 ncRNAs that are listed shows that the majority belong to lncRNAs, many of which are novel or are labeled as antisense (AS) genes. Other prominent biotypes include pseudogenes (PG), both processed and unprocessed, reflecting their origins as duplicated genes or those generated via retro-transposition, respectively. Notably, a significant portion of PGs are transcribed, potentially playing a regulatory role for their functional gene counterparts.

Figure 4A lists the 14 highly expressed ncRNAs that include XIST, which regulates X chromosome inactivation, as well as mitochondrial genes such as 16S and 12S rRNA (ribosomal RNAs). Among the highly expressed lncRNAs are GAS5 (known as SNHG2), SNHG1, NORAD and NEAT1 which are implicated in processes like the cell cycle, cancer progression, DDR, and inflammation, respectively (35).

**Figure 4.**
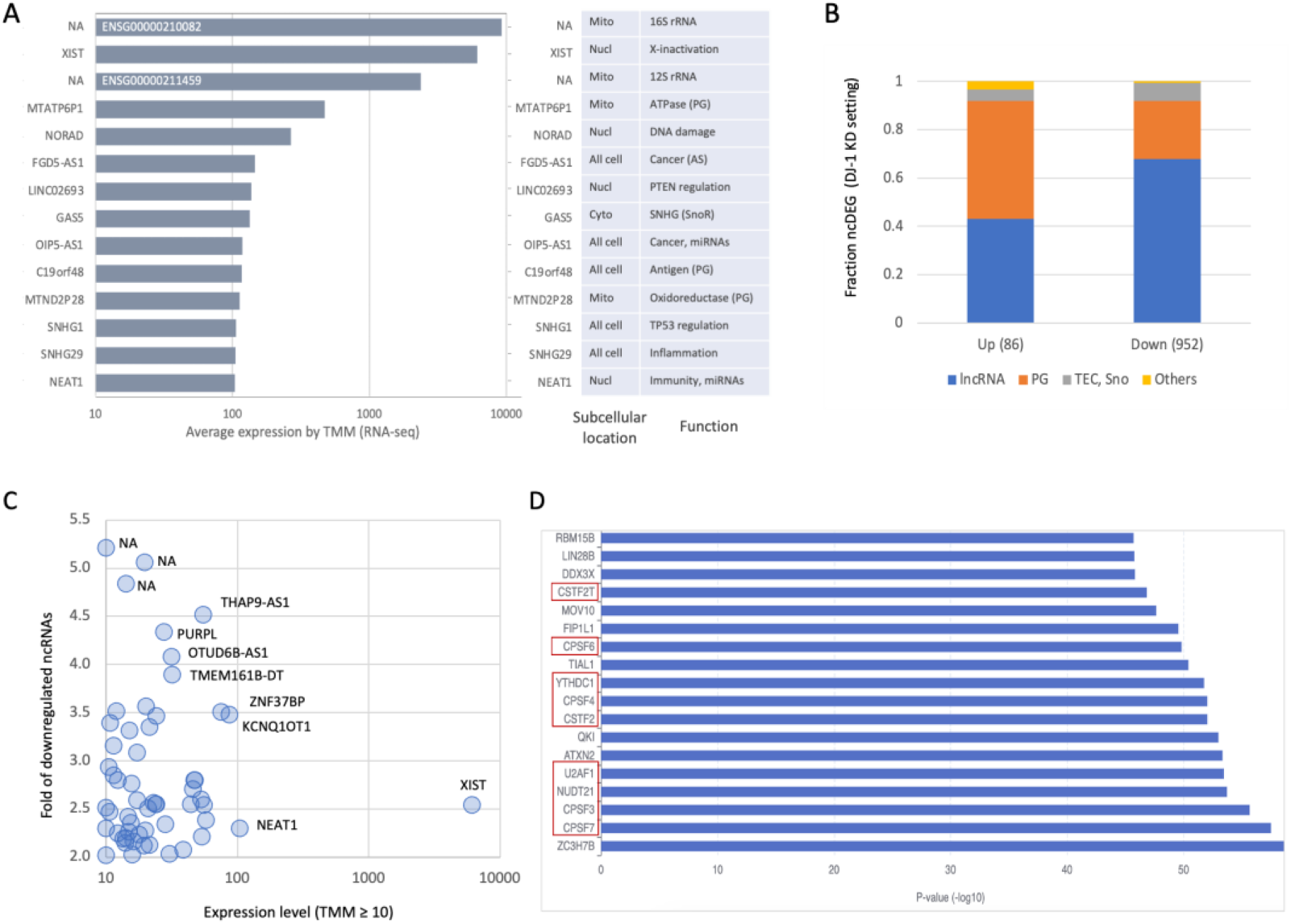
Quantitative summary of the ncRNAs by X-ray irradiation on DJ-1 KD cellular setting. **(A)** List of top expressing ncRNAs (≥100 TMM) along their subcellular localization and main function. **(B)** Partition of major biotypes of ncRNAs for the ncDEG of DJ-1 KD setting for up- and downregulated genes. Only genes with FC ≥|2| are included to secure robustness. **(C)** Scatter plot of the downregulated ncDEG (log scale, with TMM ≥10, on x-axis) and the inverse fold change in the expression level (4 means the expressing level is 0.25 of the matched control cells, y-axis). Several of the genes are marked by the gene symbol. **(D)** Statistically significant candidate coding genes according for most significant 100 ncDEGs analyzed for the RNA-protein interaction category in LncSEA 2.0. Red frames are set of proteins working together for pre-mRNA processing. List of the protein-interaction predicting results is accessible in Supplementary **Table S3**.

### X-ray irradiation on siRNA DJ-1 KD resulted in a global dysregulation of ncRNA transcription

The composition of all 1038 ncDEG that were associated with KD in the DJ-1 setting (with a strict FC threshold of ≥|2|) is shown in Figure 4B. The fraction of PGs from all identified ncRNAs in the DJ-1 KD setting is 2.5 folds higher among the upregulated ncDEG (Figure 4B). It is possible that the transcribed PGs change the execution of the transcriptional machinery. Inspecting the PGs that were downregulated, we identified about 30% of the sequences to be related to ribosomal and mitochondrial proteins, while the vast majority (70%) were uncharacterized ncRNAs (Supplementary **Table S1**). The accepted notion that most ncRNAs are unstable (i.e., fast degrading); we focused on downregulated ncDEG whose baseline expression is modest (≥10 TMM) and their downregulation was substantial (≥2 folds). Figure 4C indicated that among this set (total 56 ncRNAs), several lncRNAs belong to the most stable ncRNAs (e.g., XIST, NEAT1). Many of the other ncRNAs have been studied in the context of cancer (e.g., THAP9-AS1, PURPL, OTUD6B-AS1), regulation of transcription (e.g., KCNQ1OT1), and chromatin structure.

We used a compilation of several tools (see Methods) to create a ranked list of proteins that are more likely to be affected by ncDEG (Figure 4D). Interestingly, the candidate proteins that are likely to be affected by the change in expression of lncRNA highlighted a set of RNA-binding proteins that coordinately act on 3’ RNA processing and pre-RNA splicing (Figure 4D). Specifically, NUDT21 is a component of the 3’ RNA cleavage and polyadenylation processing. A rich set of CSTF genes (Figure 4D, red frames) act as activators of the pre-mRNA 3’-end cleavage and polyadenylation processing required for the maturation of mRNAs. Many of these nuclear RNA-binding proteins act on the 3’ end cleavage on mRNAs and consequently indirectly also impact miRNA biogenesis (e.g., ZC3H7B). For details on the enrichment summary, see **Supplemental Table S3**.

### A tight control on ncRNA transcription landscape following X-ray irradiation in DJ-1 OX cells

The overall number of ncDEG in the DJ-1 OX setting was <10% of the ncDEG associated with DJ-1 KD (Figure 2C). By testing the reliability of the expression levels of the ncRNAs in DJ-1 OX setting we showed that quantitation was extremely stable. The two data sets (DJ-1 OX and DJ-1 OX X-ray) are strongly correlated with R^2^ of 0.994 (Figure 5A, left). Notably, such strong correlation holds for the lowest set of ncRNAs (R^2^ 0.945, Figure 5A, right), confirming the tight regulation of their expression (Figure 5B). We concluded that despite the very low level of expression, the data is reliable across all ranges.

**Figure 5.**
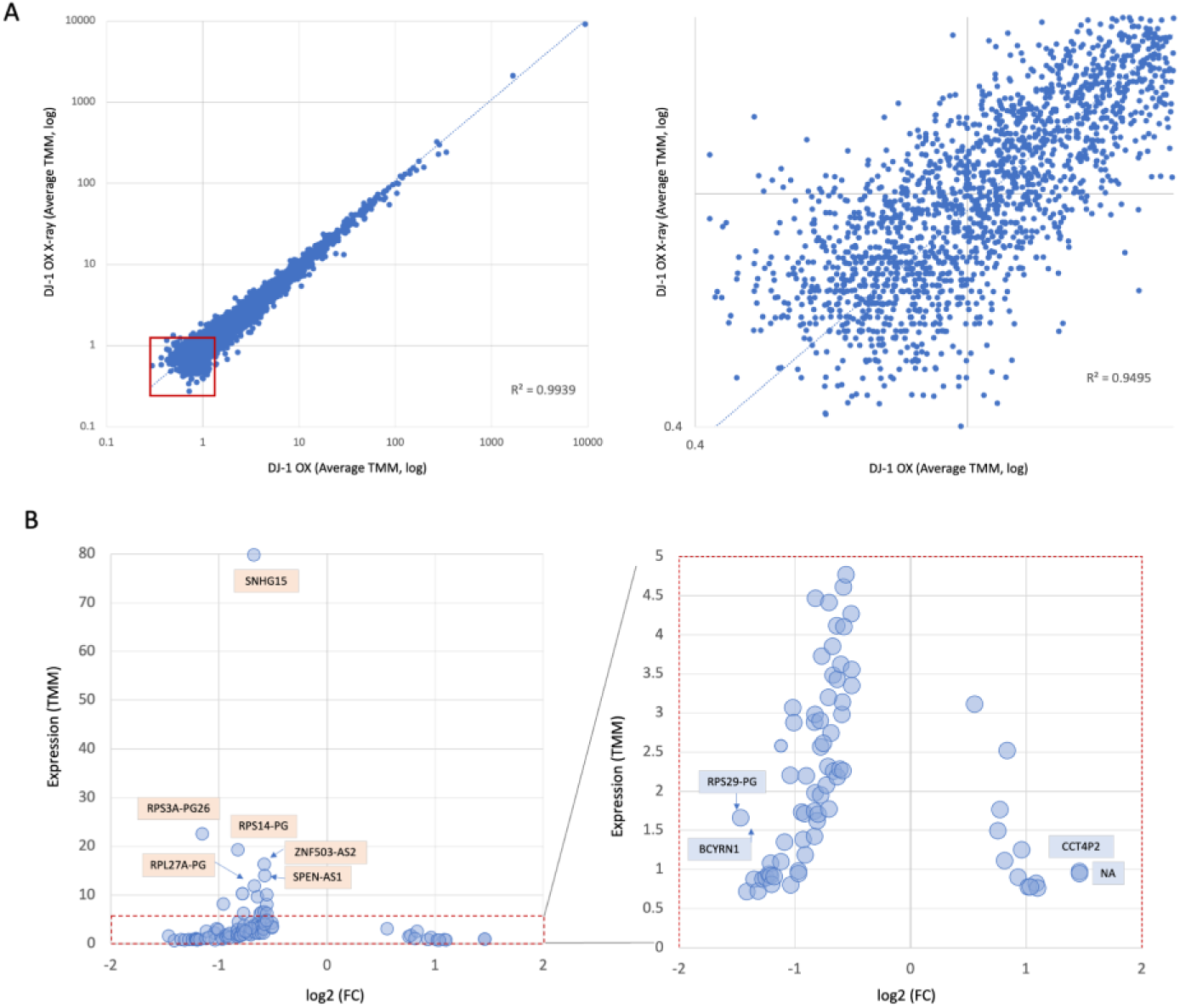
Quantitative view on ncRNA of DJ-1 OX setting. **(A)** Scatter plot for the expression level (TMM, log) of the DJ-1 OX and DJ-1 OX post X-ray at all expression range (left). A zoom in at the lowest expression set (0.4 to 2 TPM, right). **(B)** ncDEG of the DJ-1 OX setting by the log2(FC) and the expression level (TMM, left). Scatter plot of all ncDEG with fold increase ≥|1.4|, versus the amounts by TMM (total 96 genes). The ncDEG with their names (orange background) are expressed at a minimal level of ≥10 TMM (left). Zoom-in of the ncDEG expression <5 TMM (right, dotted red frame). The labelled genes (light blue) exhibit highest fold change in expression.

Testing the 96 ncDEG from the X-ray exposure cells in OX DJ-1 (Figure 5B) showed that most ncDEG were downregulated and their level of expression was very low (<5 TMM). Focusing on the lower expressing ncDEG (Figure 5B, right) confirmed that the absolute fold change is subtle for all ncDEG (fold change of 1.4 to 2.8 folds). Several ncRNAs with maximal log2(FC) were labelled (BCYRN1, RSP29- PG and CCT4P2; Figure 5B, right). Interestingly, following X-ray irradiation BCYRN1 (also called BC200) that was implicated in cell survival and migration retained 40% of its baseline level. Moreover, BCYRN1 negatively regulated translation via its impact on mRNA stability, translation, and splicing. BCYRN1 directly binds several RNA binding proteins (e.g., PABPC1) of ribosomal subunits, and polysomal RNAs (36). A common theme among the listed ncDEG is their immediate impact of general transcription and chromatin structure. We inspected the function of upregulated genes for the DJ-1 OX cellular setting. Among them we identified lncRNAs that impact chromatic structure. Example is a lncRNA that complements YEATS2, a chromatin reader of the histone acetyltransferase complex. We concluded that after X-ray irradiation on the background of DJ-1 OX the shifting in overall transcription is robust and reproducible.

### Potential functions of ncRNAs as antisense in response to X-ray irradiation

Figure 6 presents a few candidates of the ncDEG identified in the DJ-1 OX settings with the following functions: (i) antisense (AS) that impact transcriptional regulation; (ii) members of the SNHG family that are host genes for snoRNAs; (iii) lncRNA-protein candidates. Figure 6A highlights how the lncRNA SMARCA5-AS1 might lock SMARCA5 transcription by competing with the 5’ exon of the transcript. SMARCA5 (SNF2H) is an essential ATPase involved in chromatin remodeling and plays a key role in the DDR (37). By modifying chromatin structure, it facilitates access to damaged DNA for major repair mechanisms, while SMARCA5 depletion led to increased sensitivity to ionizing radiation and impaired DSB repair (38). The CHUK-DT is an upregulated lncRNA that complements the CWF19 family member (Figure 6B). CHUK-DT consists of a diverse collection of alternatively spliced ncRNAs, several of which are transcribed in the opposite direction, potentially interfering with transcription. CWF19L1 (CWF19- like cell cycle control factor 1) overlaps with the exon sequence (Figure 6B). Overexpression of CWF19L1 has been shown to inhibit the G1/S phase transition by suppressing cell cycle proteins, such as CDK4 and CDK6 (39). Figure 6C shows the MORF4L2-AS1 which may act as antisense for MORF4L2, its downregulation could result in increase in the amount of MORF4L2, a gene involved in positive regulation of transcription by RNA PolII, and associated with heterochromatin assembly and histone modification. Figure 6D illustrates the potential transcriptional interference for MYNN gene is involved in immune responses and inflammation and acts in the control of gene expression.

**Figure 6.**
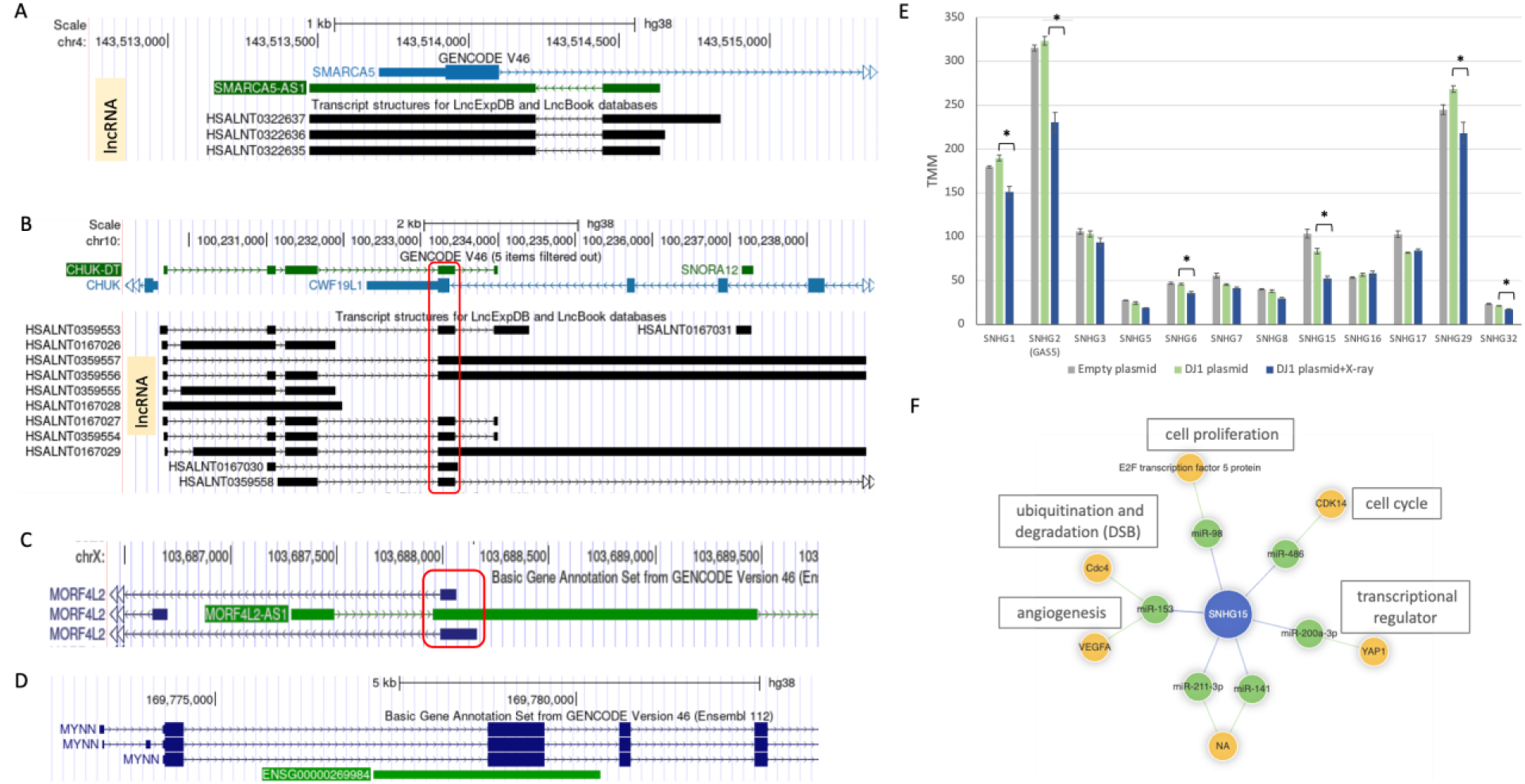
Functional candidates of ncDEG in the DJ-1 OX following X-ray. Genome browser view according to GENCODE V46. Examples of lncRNAs as antisense (AS) and the overlap on exon boundaries are shown. **(A)** Transcription overlapping of SMARCA5-AS1 and SMARCA5. The transcripts overlap the overlap the 3’-exon by complementation. **(B)** Transcription overlapping of CHUK-DT. In red frame, the overlap in exon boundaries with CWF19L1 in the complementary strand. **(C)** Transcription overlapping of MORF42L2-AS1 and MORF42L2. In red frame, the overlap of the first exon with the complementary strand. **(D)** Transcription overlapping of ENSG00000269984 and several versions of MYNN. **(E)** Relative expression of 14 members of SHNGs with ≥20 TMM. The majority of the SHNGs were downregulated 6 hours after X-ray exposure. There are 6 of the genes in which the p values (t-test) were significant (marked by asterisk). The results from empty vector (gray) show no statistical significance relative of DJ-1 OX cells. Details are available in Supplementary **Table S4**. **(F)** The interactome of SNHG15 (based on lncSEA 2.0) lists the miRNA direct interactors and their key protein targets that potentially impact the listed cellular processes.

### Suppression of SNHG family members following X-ray irradiation in DJ-1 OX cells

Among the highest expressing ncRNAs whose function was previously studied (≥100 TMM, Figure 4A), there are several genes that belong to the SHNG family (Note that GAS5 is also called SNHG2). We confirmed the statistical stability of these examples relative to controls (Supplemental **Figure S6**). GAS5 was downregulated by X-ray in cells of DJ-1 OX. However, its expression level was unchanged between cells with DJ-1 OX and cells that were introduced by an empty plasmid (Supplemental **Figure S6**). From functional perspective, GAS5 was shown to be localized to the mitochondria where it acts to maintain energy homeostasis (40). All listed examples (Supplementary **Figure S6**) displayed a similar trend. From a functional perspective, many of the identified ncDEG are also regulated by miRNAs. For example, GAS5 was negatively regulated by miR-21 (41). MINCR whose main role is in facilitating the expression of MYC-target genes that are crucial for cell cycle progression, is modulated by miR-146b- 5p in the context of NF-kB and inflammation (42).

We focused on lncRNAs of the family SNHGs (Small nucleolar RNA host genes, Figure 6E). These genes serve as hosts to snoRNAs (small nucleolar RNAs), which play key roles in rRNA modification and ribosome biogenesis. Many SNHGs also function as cytoplasmic lncRNAs with regulatory roles in gene expression, cell cycle, and cancer progression (35). We identified total of 21 SNHG genes that account for ∼8% of the total ncRNA expression in cells. Among the genes that were significantly downregulated, SNHG accounts for ∼24% of all ncRNA genes for the DJ-1 OX setting (Supplementary **Table S1**). Figure 6E shows the results of 12 SNHG family members (filtered by minimal expression level, ≥20 TMM), among them the expression of 6 were significantly suppressed. For example, SNHG15 expression level was reduced to 62% of its baseline. GAS5 (SNHG2), SNHG29, SNHG1 the downregulation following X-ray on the background of DJ-1 OX cells retailed 71%, 81% and 80%, of their level prior to the irradiation, respectively. The highly expressed SNHG15 (Figure 6F) is involved in regulating inflammatory markers (e.g., p65, TNF-α) by binding to miR-141 thus affecting the SIRT1 axis (43). Due to the high levels of SNHG15 in the cytoplasm, it is likely that such miRNA availability can be shifted. Across different cells, it was shown that gene expression regulation can be shifted by SNHGs via sponging of miRNAs (44,45). Figure 6F shows the interactome of SNHG15. Direct interaction of SNHG15 with miRNAs (e.g., miR-486, miR-200-3p), impact multiple cell responses including proliferation and cell cycle. This interactome view is in agreement with the observation that suppression of SNHG15 leads to cell cycle arrest in cancer context. We explain the coordinated suppression of SNHGs following exposure to X-ray by direct and indirect effects of DJ-1.

### Cells with overexpressed or depleted DJ-1 resulted in an inverse expression trend

We tested whether there is a ncDEGs which is a molecular hub that response to X-ray in the settings of DJ-1 OX and DJ-1 KD. A Venn diagram for the ncDEGs for N.T., DJ-1 KD, and DJ-1 OX revealed that the overlap of ncRNA is negligible (Supplementary **Figure S7**). The set of ncDEG that specifies the X-ray response in DJ-1 KD were almost entirely unique (98.6%), and there were no ncRNAs that are shared across the cellular settings (Supplementary **Figure S7**). This is in agreement with the observation regrading codDEG in which genes specified almost exclusively each cellular setting. Nevertheless, there were 12 shared ncRNAs (Supplementary **Figure S7**) that were assigned as ncDEG in DJ-1 OX and DJ-1 KD setting.

**Table 1** shows that most ncDEGs common to both DJ-1 OX and DJ-1 KD settings are classified as antisense lncRNAs (L-AS) or pseudogenes (PG). To investigate whether processed PG might function as transcriptional regulators via hybridization, we conducted a BLAST search against the entire transcriptome database, identifying genes with high-confidence matches to coding transcripts (BLAST E-score <e-20). We found that most PGs can successfully hybridize either to a similar PGs (if present) or to their gene of origin (**Table 1**). The roles of L-AS ncDEGs may influence mRNA dynamics (e.g., INTS6- AS RNA 1), chromatin structure (e.g., ATXN1-AS1), transcription regulation (e.g., MYNN antisense), and the chromatin-transcription axis (e.g., SMARCA5-AS1; Figure 6A). **Table 1** also shows that the trend for upregulation (U) and downregulation (D) was inversed between OX and KD settings for 8 of the 12 genes (**Table 1**, bold). All five genes listed (**Table 1**) that were upregulated in DJ-1 OX setting showed an inverse expression trend in DJ-1 KD setting. These genes with inverse expression trend may be considered to have a direct link to DJ-1 cellular status.

**Table 1.** ncDEG shared between DJ-1 OX and DJ-1 KD cellular settings.

| Ensembl | Gene name | Type<sup>a</sup> | Gene Description | # of exons | Size (nt) | Hyb<sup>b</sup> | Trend OX | Trend KD |
| --- | --- | --- | --- | --- | --- | --- | --- | --- |
| ENSG00000225569 | CCT4P2 | PG | chaperonin w TCP1 subunit 4 PG2 | 3 | 1548 | S | U | D |
| ENSG00000234084 | NA | Lnc | Novel | 3 | 577 | C | U | D |
| ENSG00000242600 | NA | TPG | mannose binding lectin 1, PG | 4 | 758 | - | U | D |
| ENSG00000245112 | SMARCA5-AS1 | L-AS | SMARCA5 AS RNA 1 | 2 | 1149 | C | U | D |
| ENSG00000269984 | NA | L-AS | novel, AS of MYNN | 2 | 3143 | C | U | D |
| ENSG00000198221 | NA | L-DT* | AFDN DT | 2 | 4095 | S | D | D |
| ENSG00000218426 | RPL27AP6 | PG | 60S ribosomal protein L27a PG | 1 | 447 | S | D | U |
| ENSG00000226415 | NA | PG | triosephosphate isomerase 1 PG1 | 1 | 750 | S | D | U |
| ENSG00000229931 | NA | L-AS | ATXN1 AS RNA 1 | 3 | 1913 | - | D | D |
| ENSG00000231154 | NA | L-AS | MORF4L2 AS RNA 1 | 3 | 2086 | C | D | D |
| ENSG00000236778 | INTS6-AS1 | L-AS* | INTS6 AS RNA 1 | 2 | 4305 | - | D | D |
| ENSG00000276168 | RN7SL1 | mis. | RNA of signal recognition particle | 1 | 299 | - | D | U |

<sup>a</sup>TPG, transcribed pseudogene; Lnc, L-, long noncoding RNA; AS, antisense; mis., miscellaneous; \*, multiple transcripts. <sup>b</sup>Hyb, hybridization by BLAST E-value <1e-20. Match to same strand direction (S), and the complementary strand (C). <sup>b</sup>Size, in case of multiple transcripts, the longest/ representative one is reported.

### Enrichment of cell cycle arrest and chromatin functional classes under varying DJ-1 expression levels

Most reported lncRNAs (Supplementary **Table S1**) are low expressing and poorly annotated. Therefore, assigning possible functions to any specific lncRNA is challenging. Instead, we assessed the potential role of the ncRNAs as a set. To this end, we applied an enrichment scheme (based on LncSEA 2.0) for the collection of ncRNAs that were downregulated by X-ray in DJ-1 OX settings (55 of the 81 most significant ncRNAs were uniquely mapped; see Methods).

Several classes of chromatin regulators and histone modifications were enriched among the downregulated genes from DJ-1 OX cellular setting (**Table 2**). FAM66C, FLJ37453 and SNHG10 were identified by ChIP-seq to act as chromatin regulation (adjusted p-value 3.5e-04). It is corroborated with the observation that 50% of all the downregulated mapped lncRNAs (25 genes) were known to bind H3K4me2-3 (Adj. p-value: 9e-23; **Table 2**), another histone marker is associated with phosphorylated of H3 at Thr11 (H3T11P; Adj. p-value 5e-16). Such enrichment results suggest that many of the downregulated lncRNAs impact gene expression regulation through binding to histone epigenetic marks.

**Table 2.** Functional enrichments of downregulated genes from DJ-1 OX cellular setting by LncSEA2.0.

| Class (LncSEA) | Set | # (query) | Size of set | P-value | Adj. p-value | Jaccard | Simpson <sup>a</sup> |
| --- | --- | --- | --- | --- | --- | --- | --- |
| RNA Histone Modification | H3K4me2-3 | 25 | 1802 | 1.80E-23 | 9.00E-23 | 0.014 | 0.455 |
|  | H3T11P | 20 | 1863 | 1.84E-16 | 4.60E-16 | 0.011 | 0.364 |
|  | H3K122ac | 20 | 1991 | 6.48E-16 | 1.08E-15 | 0.010 | 0.364 |
|  | H2AFX | 6 | 945 | 2.59E-04 | 3.24E-04 | 0.006 | 0.109 |
| Chromatin Regulators | NCOA1 | 3 | 48 | 1.32E-05 | 3.99E-03 | 0.030 | 0.063 |
| Experimental Validated Function | cancer progression | 7 | 307 | 1.68E-08 | 9.07E-07 | 0.020 | 0.127 |
|  | cell metastasis | 6 | 264 | 1.93E-07 | 6.94E-06 | 0.019 | 0.109 |
|  | cell cycle | 5 | 143 | 2.57E-07 | 6.94E-06 | 0.026 | 0.091 |
|  | proliferation | 8 | 838 | 1.16E-06 | 2.51E-05 | 0.009 | 0.145 |
| RNA Protein Interaction | DDX3X | 26 | 961 | 6.65E-32 | 6.69E-28 | 0.026 | 0.473 |
|  | ATXN2 | 26 | 1032 | 4.20E-31 | 2.11E-27 | 0.025 | 0.473 |
|  | CSTF2T | 26 | 1328 | 2.75E-28 | 9.23E-25 | 0.019 | 0.473 |

<sup>a</sup>Simpson assesses the diversity in the functional roles of lncRNAs in the database.

**Table 2** also identified numerous experimental validated sets (based on LncTarD2.0) for cell growth, cancer progression, cell cycle and more (Adj. p-values <2e-05). The 8 ncDEG that are listed for cell proliferation are BCYRN1, FOXCUT, HOXA11-AS, LINC01535, MINCR, MNX1-AS1, SNHG10 and SNHG15.

By considering the downregulated lncRNAs as a group, serval target genes emerged for the class of RNA-protein interaction. Examples are DDX3X, ATXN2 and CSTF2T for which 26 of the 55 mapped genes were involved (Adj. p-value ranges from 6.6e-32 to 2.8e-28). These are versatile proteins involved in various aspects of RNA metabolism, cell cycle, innate immunity, and apoptosis and translation. DDX3X was also directly involve in DNA damage response, indicating its role in genome stability, while ATXN2 and CSTF2T are proteins that act on the mRNA molecule in the cytoplasm. ATXN2 assembles with polysomes and interacts with the poly(A)-binding proteins (PABPC1) to dictate translation initiation and mRNA decay regulation. CSTF2T was predicted to be involved in pre-mRNA cleavage required for polyadenylation. We concluded that upon exposure to X-ray, the genome stability is affected and, through lncRNA alternation enzymes that function in RNA metabolism and mRNA stability, shift to cope with the new cellular conditions.

## Discussion

In this study, we assess how cells respond to X-ray irradiation at varying levels of DJ-1 by examining the differential transcriptome landscape. Most studies on DJ-1 focus on its fundamental role in sensing and responding to oxidative stress. The modified DJ-1 protein shuttles from the cytosol to the mitochondria and then to the nucleus, impacting the antioxidation response (46). The cellular protective role of DJ-1 is attributed to its function in the mitochondria. DJ-1 increases the expression of uncoupling proteins, suppressing ROS production while stabilizing interaction with mitochondrial Bcl-xL and consequently inhibiting apoptosis activation (47). At the transcriptional level, DJ-1 stabilizes Nrf2, facilitating its translocation into the nucleus to act as a major transcription factor for antioxidant genes (5).

The notion that DJ-1 might activate alternative mechanisms driving cells toward pathological states has also been proposed (1,48). We show that DJ-1 impacts cell decisions and the transcriptomic landscape following genotoxic stress, emphasizing its nuclear role. Recent studies suggest that DJ-1 regulates DSB repair and the DDR in the context of PD (49). While irradiation may cause transient oxidation, we confirm that the tested cells activate DDR without changes in apoptotic levels (Supplementary Figure S1). The treatment with X- rays causes DSBs that are effectively addressed by γ- H2Ax to initiate DDR. Specifically, the phosphorylation of histone variant H2Ax is essential for HR and NHEJ DNA repair pathways (50). The irradiated cells recovered successfully from the X-ray insult with minimal signs of cell death. To isolate the impact of X-ray, we used matched controls with predetermined DJ-1 expression levels. Specifically, cells with non-specific siRNA (siRNA RULC) served as controls for DJ-1 KD settings, while empty plasmid-transfected cells served as controls for DJ-1 OX. To address the importance of DJ-1 at the protein level, we showed that the DJ-1 OX of the DJ-1 transcript is validated at the protein level. A similar effect was confirmed in the DJ-1 KD with strong suppression of DJ-1 protein levels (Supplementary Figure S2). Notably, unlike the robust antiviral response induced by siRNA manipulation (33), no antiviral-like response was observed 6 hours post X-ray exposure. We did not study the kinetics of DNA recovery. However, it is expected to vary by tissue and cell type (51).

A key finding in the DJ-1 OX setting is the downregulation of ribosomal genes following X-ray irradiation, including mitochondrial ribosomal genes (Figure 3F). Ribosome assembly in humans requires coordinated RNA polymerase activities for rRNA processing, RNA modifications, and assembly. As ribosome biogenesis consumes significant cellular resources, its regulation is vital for homeostasis.

The observation that dozens of ribosomal transcripts were downregulated by X-ray in the DJ-1 OX setting aligns with the lability of the unassembled ribosomal genes, triggering ribosomal stress (52). Altered mitochondrial function and nutrient deficiency may also contribute to the observed downregulation. The connection between ribosome biogenesis and cell growth highlights p53 signaling’s involvement in stress responses (53,54). The interaction of DJ-1 with p53 has been studied in PD and cancer (5). Following X-ray exposure and DSB formation, p53 initiates cell cycle arrest and DNA repair, with its activity strongly regulated by MDM2 (55). Disruptions in ribosome biogenesis (i.e., nucleolar stress), as observed in DJ-1 OX cells, can trigger the MDM2-p53 stress response. Interestingly, ribosomal proteins such as RPL5, RPL11, and RPL23 can directly bind MDM2, thereby stabilizing p53 (56). The impact of X-ray in the DJ-1 OX and DJ-1 KD settings was evaluated through inspecting the codDEG, particularly lncRNAs. In general, our results support the global shift in cellular homeostasis. However, we noted that ncDEG also play a fundamental role in the observed transcriptome landscape. Approximately 4,200 ncRNAs were detected in DJ-1 KD cells, of which 25% were identified as ncDEG (log2(FC) >|1|). This significant number likely reflects reduced cell resilience under nearly complete DJ-1 depletion (33). Several observations support the notion of cellular fragility. Specifically, more than 1,000 lncRNAs changed within 6 hours of X-ray exposure, with 92% showing reduced expression. Moreover, the ncDEG in DJ-1 KD and DJ-1 OX cells represented nearly disjoint sets.

A surprising observation is the strong coordinated response of SHNG transcripts to X-ray irradiation in DJ-1 OX cells. These genes are associated with cancer progression (57). Among the 12 of 21 family members with high expression (≥20 TMM), half were also significantly suppressed (Figure 6E). SnoRNAs, which are primarily responsible for rRNA modifications, are hosted within SHNG gene introns. Additionally, SnoRNAs are located within ribosomal protein genes (58). The downregulation of SHNG genes and ribosomal proteins suggests ribosome destabilization through direct effects on rRNAs. Moreover, SHNG genes such as SNHG15 affect miRNA sponging, which acts in post-transcriptional regulation. For instance, SNHG15 (a host for SNORA9) influences miRNA activity beyond SnoRNA production (Figure 6F). The DJ-1 response to X-ray may also indirectly impact miRNA profiling by influencing damage-induced RNAs (diRNAs) at DSB sites (59), which recruit Argonaute 2 (AGO2) and affect miRNA biogenesis. The miRNA-mediated regulation influences cell survival pathways beyond direct effects on DNA repair (60).

Several lncRNAs upregulated in DJ-1 OX cells post-X-ray are linked to transcription. For instance, PAIP1P1, derived from the PAIP1 gene, may influence translational initiation and biosynthesis. ENSG00000229447, a pseudogene related to GTF2A2, ensures accuracy in transcription initiation. Other ncDEG are crucial to homeostatic functions, such as TPI1P2 (glycolysis), CCT4P2 (protein folding), and BCLAF1P2 (mRNA stability). lncRNAs also play a role in governing the complexity of stress responses and shifts in homeostasis, impacting pathways like p53 activation (61,62). SMARCA5-AS1, involved in chromatin remodeling and maintaining genomic integrity (63), dynamically responds to DJ-1 levels and X-ray irradiation (Table 1). Its altered expression suggests regulatory roles in chromatin architecture and gene expression.

DJ-1 OX cells exhibit restricted and robust transcriptional changes post-X-ray exposure. Elevated DJ-1 levels in cancer samples correlate with advanced stages and poor prognosis (64). DJ-1’s role in coordinating signaling pathways, especially in cancer progression, remains elusive (48). For example, DJ-1 increases cyclin-D1 expression in colorectal cancer, driving cell cycle progression, and suppresses p53-mediated apoptosis under genotoxic stress (65). In contrast, DJ-1 KD cells show widespread transcriptional alterations mediated by lncRNAs, affecting the cell cycle, proliferation, mRNA maturation, miRNA stability, and RNA-dependent nuclear functions.

## Abbreviations

AGO2: Argonaute 2
DDR: DNA damage response
DEG: differentially expressed genes
DSB: double-strand break
HR: homologous recombination
KD: knockdown
N.T.: not treated
ncRNA: non-coding RNA
NHEJ: non-homologous end joining
Nt: nucleotides
OX: overexpression
PCA: principal component analysis
PD: Parkinson’s disease protein-protein interaction (PPI)
ROS: reactive oxygen species
TF: transcription factor
TMM: trimmed mean of the M-values normalization

## Supplementary Data

Figure S1-S7.

**Table S1.** The summary of all DEGs based on the 18 RNA-seq samples used in this study. The 18 samples are used to provide p-value [FDR], the DEG log_2_FC, biotypes and transcript names. The cellular settings included are: (i) [N.T. X-Ray] vs [NT], (ii) [DJ-1 OX X-Ray] vs [DJ-1 OX], and (iii) [DJ-1 KD X-Ray] vs [DJ-1 KD].

**Table S2.** Partition to up and downregulated codDEG for the KD setting. Source for Figure 3E and Supplementary Figure S4.

**Table S3.** The enrichment results for the downregulated ncDEG of DJ-1 KD [X-Ray] vs [DJ-1 KD]. The input were 100 transcripts, 85 were successfully mapped by the ID convertor of lncSEAs 2.0. The enrichment refer only to the RNA-protein interaction category. The findings are sorted and filtered by a p-value [FDR] of <1e-09. Source for Figure 4D.

**Table S4.** Statistical results for the members of the SHNG set. Source for Figure 6E.

## Author contributions

K.Z. led the project from the experimental design to the analysis and visualization. M.L. and M.G. served in mentoring. optimized the X-ray experiments and validation by antibodies. T.E. contributed in preparation for RNA extraction and sequencing libraries for RNA-seq sequencing. H.O.Z. conducted the

Western blots. M.L. wrote the initial manuscript, with all coauthors read, commented and contributed to the final version.

## Acknowledgments

We thank the members of the Linial’s and Goldberg’s labs for useful suggestions and fruitful discussions. We thank the Genomic and Sequencing Center of the Hebrew University for their constant support. We thank the system team of the Computer Science and Engineering, and the Computer Services at the Hebrew University for their support. We thank the Clore Israel Foundation for their Scholar program and the fellowship granted to K.Z.

## Funding

This study was supported by the ISF grant 2753/20 (M.L.), a grant (2023) by the National Alopecia Areata Foundation (NAAF) on Genetics of alopecia areata (M.L.) and a fellowship from Center for Interdisciplinary Data Science Research (3035000440).

## Data availability

All processed and normalized data produced in this study are available in Supplementary Tables. The raw data was submitted to ArrayExpress Accession E-MTAB-14761.

## Conflict of interest

The authors declare that they have no conflict of interest.

## References

1. Biosa, A., Sandrelli, F., Beltramini, M., Greggio, E., Bubacco, L. and Bisaglia, M. (2017) Recent findings on the physiological function of DJ-1: Beyond Parkinson’s disease. Neurobiology of disease, 108, 65–72.

2. Riepe, C., Zelin, E., Frankino, P.A., Meacham, Z.A., Fernandez, S.G., Ingolia, N.T. and Corn, J.E. (2022) Double stranded DNA breaks and genome editing trigger loss of ribosomal protein RPS27A. FEBS J, 289, 3101–3114.

3. Zhang, L., Wang, J., Wang, J., Yang, B., He, Q. and Weng, Q. (2020) Role of DJ-1 in immune and inflammatory diseases. Frontiers in immunology, 11, 529121.

4. Ariga, H., Takahashi-Niki, K., Kato, I., Maita, H., Niki, T. and Iguchi-Ariga, S.M. (2013) Neuroprotective function of DJ-1 in Parkinson’s disease. Oxid Med Cell Longev, 2013, 683920.

5. Dolgacheva, L.P., Berezhnov, A.V., Fedotova, E.I., Zinchenko, V.P. and Abramov, A.Y. (2019) Role of DJ-1 in the mechanism of pathogenesis of Parkinson’s disease. Journal of Bioenergetics and Biomembranes, 51, 175–188.

6. Kahle, P.J., Waak, J. and Gasser, T. (2009) DJ-1 and prevention of oxidative stress in Parkinson’s disease and other age-related disorders. Free Radical Biology and Medicine, 47, 1354–1361.

7. Inberg, A. and Linial, M. (2010) Protection of pancreatic β-cells from various stress conditions is mediated by DJ-1. Journal of Biological Chemistry, 285, 25686–25698.

8. Kim, J.-M., Jang, H.-J., Choi, S.Y., Park, S.-A., Kim, I.S., Yang, Y.R., Lee, Y.H., Ryu, S.H. and Suh, P.-G. (2014) DJ-1 contributes to adipogenesis and obesity-induced inflammation. Scientific reports, 4, 4805.

9. Shi, S.Y., Lu, S.-Y., Sivasubramaniyam, T., Revelo, X.S., Cai, E.P., Luk, C.T., Schroer, S.A., Patel, P., Kim, R.H. and Bombardier, E. (2015) DJ-1 links muscle ROS production with metabolic reprogramming and systemic energy homeostasis in mice. Nature communications, 6, 7415.

10. Huang, M. and Chen, S. (2021) DJ-1 in neurodegenerative diseases: Pathogenesis and clinical application. Progress in Neurobiology, 204, 102114.

11. Liu, Y., Ma, X., Fujioka, H., Liu, J., Chen, S. and Zhu, X. (2019) DJ-1 regulates the integrity and function of ER-mitochondria association through interaction with IP3R3-Grp75-VDAC1. Proceedings of the National Academy of Sciences, 116, 25322–25328.

12. Dash, B.K., Urano, Y., Saito, Y. and Noguchi, N. (2022) Redox-sensitive DJ-1 protein: an insight into physiological roles, secretion, and therapeutic target. Redox Experimental Medicine, 2022, R96–R115.

13. Wilson, M.A. (2011) The role of cysteine oxidation in DJ-1 function and dysfunction. Antioxidants & redox signaling, 15, 111–122.

14. Girotto, S., Cendron, L., Bisaglia, M., Tessari, I., Mammi, S., Zanotti, G. and Bubacco, L. (2014) DJ-1 is a copper chaperone acting on SOD1 activation. Journal of Biological Chemistry, 289, 10887–10899.

15. Oh, S.E. and Mouradian, M.M. (2018) Cytoprotective mechanisms of DJ-1 against oxidative stress through modulating ERK1/2 and ASK1 signal transduction. Redox biology, 14, 211–217.

16. Gan, L., Johnson, D.A. and Johnson, J.A. (2010) Keap1-Nrf2 activation in the presence and absence of DJ-1. European Journal of Neuroscience, 31, 967–977.

17. Chakkittukandiyil, A., Sajini, D.V., Karuppaiah, A. and Selvaraj, D. (2022) The principal molecular mechanisms behind the activation of Keap1/Nrf2/ARE pathway leading to neuroprotective action in Parkinson’s disease. Neurochemistry International, 156, 105325.

18. Kim, R.H., Peters, M., Jang, Y., Shi, W., Pintilie, M., Fletcher, G.C., DeLuca, C., Liepa, J., Zhou, L., Snow, B. et al. (2005) DJ-1, a novel regulator of the tumor suppressor PTEN. Cancer Cell, 7, 263–273.

19. Singh, M.H., Brooke, S.M., Zemlyak, I. and Sapolsky, R.M. (2010) Evidence for caspase effects on release of cytochrome c and AIF in a model of ischemia in cortical neurons. Neurosci Lett, 469, 179–183.

20. Jin, W. (2020) Novel insights into PARK7 (DJ-1), a potential anti-cancer therapeutic target, and implications for cancer progression. Journal of Clinical Medicine, 9, 1256.

21. Adjemian, S., Oltean, T., Martens, S., Wiernicki, B., Goossens, V., Vanden Berghe, T., Cappe, B., Ladik, M., Riquet, F.B., Heyndrickx, L. et al. (2020) Ionizing radiation results in a mixture of cellular outcomes including mitotic catastrophe, senescence, methuosis, and iron-dependent cell death. Cell Death Dis, 11, 1003.

22. Li, P., Liu, X., Zhao, T., Li, F., Wang, Q., Zhang, P., Hirayama, R., Chen, W., Jin, X., Zheng, X. et al. (2021) Comparable radiation sensitivity in p53 wild-type and p53 deficient tumor cells associated with different cell death modalities. Cell Death Discov, 7, 184.

23. Wang, Z.X., Liu, Y., Li, Y.L., Wei, Q., Lin, R.R., Kang, R., Ruan, Y., Lin, Z.H., Xue, N.J., Zhang, B.R. et al. (2023) Nuclear DJ-1 Regulates DNA Damage Repair via the Regulation of PARP1 Activity. Int J Mol Sci, 24.

24. Sawai, H. and Domae, N. (2011) Discrimination between primary necrosis and apoptosis by necrostatin-1 in Annexin V-positive/propidium iodide-negative cells. Biochem Biophys Res Commun, 411, 569–573.

25. Rieger, A.M., Nelson, K.L., Konowalchuk, J.D. and Barreda, D.R. (2011) Modified annexin V/propidium iodide apoptosis assay for accurate assessment of cell death. J Vis Exp.

26. Zohar, K., Giladi, E., Eliyahu, T. and Linial, M. (2022) Oxidative Stress and Its Modulation by Ladostigil Alter the Expression of Abundant Long Non-Coding RNAs in SH-SY5Y Cells. Noncoding RNA, 8.

27. Azzam, T., Raskin, A., Makovitzki, A., Brem, H., Vierling, P., Lineal, M. and Domb, A.J. (2002) Cationic polysaccharides for gene delivery. Macromolecules, 35, 9947–9953.

28. Hunt, M.A., Currie, M.J., Robinson, B.A. and Dachs, G.U. (2010) Optimizing transfection of primary human umbilical vein endothelial cells using commercially available chemical transfection reagents. Journal of biomolecular techniques: JBT, 21, 66.

29. Zohar, K. and Linial, M. (2024) Knockdown of DJ-1 Resulted in a Coordinated Activation of the Innate Immune Antiviral Response in HEK293 Cell Line. Int J Mol Sci, 25.

30. Chen, J., Zhang, J., Gao, Y., Li, Y., Feng, C., Song, C., Ning, Z., Zhou, X., Zhao, J., Feng, M. et al. (2021) LncSEA: a platform for long non-coding RNA related sets and enrichment analysis. Nucleic Acids Res, 49, D969–D980.

31. Li, Z., Zhang, Y., Fang, J., Xu, Z., Zhang, H., Mao, M., Chen, Y., Zhang, L. and Pian, C. (2023) NcPath: a novel platform for visualization and enrichment analysis of human non-coding RNA and KEGG signaling pathways. Bioinformatics, 39.

32. Rogakou, E.P., Pilch, D.R., Orr, A.H., Ivanova, V.S. and Bonner, W.M. (1998) DNA double-stranded breaks induce histone H2AX phosphorylation on serine 139. Journal of biological chemistry, 273, 5858–5868.

33. Zohar, K. and Linial, M. (2024) Knockdown of DJ-1 Resulted in a Coordinated Activation of the Innate Immune Antiviral Response in HEK293 Cell Line. International Journal of Molecular Sciences, 25, 7550.

34. Schlattner, U. (2021) The Complex Functions of the NME Family-A Matter of Location and Molecular Activity. Int J Mol Sci, 22.

35. Xiao, H., Feng, X., Liu, M., Gong, H. and Zhou, X. (2023) SnoRNA and lncSNHG: Advances of nucleolar small RNA host gene transcripts in anti-tumor immunity. Front Immunol, 14, 1143980.

36. Booy, E.P., Gussakovsky, D., Choi, T. and McKenna, S.A. (2021) The noncoding RNA BC200 associates with polysomes to positively regulate mRNA translation in tumor cells. J Biol Chem, 296, 100036.

37. Toiber, D., Erdel, F., Bouazoune, K., Silberman, D.M., Zhong, L., Mulligan, P., Sebastian, C., Cosentino, C., Martinez-Pastor, B. and Giacosa, S. (2013) SIRT6 recruits SNF2H to DNA break sites, preventing genomic instability through chromatin remodeling. Molecular cell, 51, 454–468.

38. Smeenk, G., Wiegant, W.W., Marteijn, J.A., Luijsterburg, M.S., Sroczynski, N., Costelloe, T., Romeijn, R.J., Pastink, A., Mailand, N., Vermeulen, W. et al. (2013) Poly(ADP-ribosyl)ation links the chromatin remodeler SMARCA5/SNF2H to RNF168-dependent DNA damage signaling. J Cell Sci, 126, 889–903.

39. Shi, J., Lv, S., Wu, M., Wang, X., Deng, Y., Li, Y., Li, K., Zhao, H., Zhu, X. and Ye, M. (2020) HOTAIR-EZH2 inhibitor AC1Q3QWB upregulates CWF19L1 and enhances cell cycle inhibition of CDK4/6 inhibitor palbociclib in glioma. Clinical and Translational Medicine, 10, 182–198.

40. Sang, L., Ju, H.-q., Yang, Z., Ge, Q., Zhang, Z., Liu, F., Yang, L., Gong, H., Shi, C. and Qu, L. (2021) Mitochondrial long non-coding RNA GAS5 tunes TCA metabolism in response to nutrient stress. Nature metabolism, 3, 90–106.

41. Zhang, Z., Zhu, Z., Watabe, K., Zhang, X., Bai, C., Xu, M., Wu, F. and Mo, Y.Y. (2013) Negative regulation of lncRNA GAS5 by miR-21. Cell Death Differ, 20, 1558–1568.

42. Gao, W. and Zhang, Y. (2021) Depression of lncRNA MINCR antagonizes LPS-evoked acute injury and inflammatory response via miR-146b-5p and the TRAF6-NFkB signaling. Mol Med, 27, 124.

43. Kang, M., Ji, F., Sun, X., Liu, H. and Zhang, C. (2021) LncRNA SNHG15 Promotes Oxidative Stress Damage to Regulate the Occurrence and Development of Cerebral Ischemia/Reperfusion Injury by Targeting the miR-141/SIRT1 Axis. J Healthc Eng, 2021, 6577799.

44. Shuai, Y., Ma, Z., Lu, J. and Feng, J. (2020) LncRNA SNHG15: A new budding star in human cancers. Cell Prolif, 53, e12716.

45. Kong, Q. and Qiu, M. (2018) Long noncoding RNA SNHG15 promotes human breast cancer proliferation, migration and invasion by sponging miR-211-3p. Biochem Biophys Res Commun, 495, 1594–1600.

46. Zohar, K., Giladi, E., Eliyahu, T. and Linial, M. (2022) Alteration in long noncoding RNAs in response to oxidative stress and ladostigil in SH-SY5Y cells. bioRxiv, 2022.2002. 2020.481187.

47. Ren, H., Fu, K., Wang, D., Mu, C. and Wang, G. (2011) Oxidized DJ-1 interacts with the mitochondrial protein BCL-XL. Journal of Biological Chemistry, 286, 35308–35317.

48. Neves, M., Graos, M., Anjo, S.I. and Manadas, B. (2022) Modulation of signaling pathways by DJ-1: An updated overview. Redox Biol, 51, 102283.

49. Wang, Z.-X., Liu, Y., Li, Y.-L., Wei, Q., Lin, R.-R., Kang, R., Ruan, Y., Lin, Z.-H., Xue, N.-J. and Zhang, B.-R. (2023) Nuclear DJ-1 Regulates DNA Damage Repair via the Regulation of PARP1 Activity. International Journal of Molecular Sciences, 24, 8651.

50. Collins, P.L., Purman, C., Porter, S.I., Nganga, V., Saini, A., Hayer, K.E., Gurewitz, G.L., Sleckman, B.P., Bednarski, J.J., Bassing, C.H. et al. (2020) DNA double-strand breaks induce H2Ax phosphorylation domains in a contact-dependent manner. Nat Commun, 11, 3158.

51. Aryankalayil, M.J., Bylicky, M.A., Martello, S., Chopra, S., Sproull, M., May, J.M., Shankardass, A., MacMillan, L., Vanpouille-Box, C., Dalo, J. et al. (2023) Microarray analysis identifies coding and non-coding RNA markers of liver injury in whole body irradiated mice. Sci Rep, 13, 200.

52. Zhou, X., Liao, W.J., Liao, J.M., Liao, P. and Lu, H. (2015) Ribosomal proteins: functions beyond the ribosome. J Mol Cell Biol, 7, 92–104.

53. Golomb, L., Volarevic, S. and Oren, M. (2014) p53 and ribosome biogenesis stress: the essentials. FEBS letters, 588, 2571–2579.

54. Lindström, M.S., Bartek, J. and Maya-Mendoza, A. (2022) p53 at the crossroad of DNA replication and ribosome biogenesis stress pathways. Cell Death & Differentiation, 29, 972–982.

55. Moll, U.M. and Petrenko, O. (2003) The MDM2-p53 interaction. Mol Cancer Res, 1, 1001–1008.

56. Liu, Y., Deisenroth, C. and Zhang, Y. (2016) RP-MDM2-p53 Pathway: Linking Ribosomal Biogenesis and Tumor Surveillance. Trends Cancer, 2, 191–204.

57. Zimta, A.A., Tigu, A.B., Braicu, C., Stefan, C., Ionescu, C. and Berindan-Neagoe, I. (2020) An Emerging Class of Long Non-coding RNA With Oncogenic Role Arises From the snoRNA Host Genes. Front Oncol, 10, 389.

58. Huldani, H., Gandla, K., Asiri, M., Romero-Parra, R.M., Alsalamy, A., Hjazi, A., Najm, M.A.A., Fawaz, A., Hussien, B.M. and Singh, R. (2023) A comprehensive insight into the role of small nucleolar RNAs (snoRNAs) and SNHGs in human cancers. Pathol Res Pract, 249, 154679.

59. Wei, W., Ba, Z., Gao, M., Wu, Y., Ma, Y., Amiard, S., White, C.I., Rendtlew Danielsen, J.M., Yang, Y.G. and Qi, Y. (2012) A role for small RNAs in DNA double-strand break repair. Cell, 149, 101–112.

60. Kraemer, A., Anastasov, N., Angermeier, M., Winkler, K., Atkinson, M.J. and Moertl, S. (2011) MicroRNA-mediated processes are essential for the cellular radiation response. Radiat Res, 176, 575–586.

61. Nie, J., Peng, C., Pei, W., Zhu, W., Zhang, S., Cao, H., Qi, X., Tong, J. and Jiao, Y. (2015) A novel role of long non-coding RNAs in response to X-ray irradiation. Toxicology In Vitro, 30, 536–544.

62. Chen, L., Ren, P., Zhang, Y., Gong, B., Yu, D. and Sun, X. (2020) Long non-coding RNA GAS5 increases the radiosensitivity of A549 cells through interaction with the miR-21/PTEN/Akt axis. Oncol Rep, 43, 897–907.

63. Thakur, S., Cahais, V., Turkova, T., Zikmund, T., Renard, C., Stopka, T., Korenjak, M. and Zavadil, J. (2022) Chromatin Remodeler Smarca5 Is Required for Cancer-Related Processes of Primary Cell Fitness and Immortalization. Cells, 11.

64. Cao, J., Lou, S., Ying, M. and Yang, B. (2015) DJ-1 as a human oncogene and potential therapeutic target. Biochemical pharmacology, 93, 241–250.

65. Fan, J., Ren, H., Jia, N., Fei, E., Zhou, T., Jiang, P., Wu, M. and Wang, G. (2008) DJ-1 decreases Bax expression through repressing p53 transcriptional activity. Journal of Biological Chemistry, 283, 4022–4030.

